# Aging-related changes in expression and function of glutamate transporters in rat spinal cord astrocytes

**DOI:** 10.1101/2023.02.14.528576

**Authors:** Shiksha Sharan, Bhanu Prakash Tewari, Preeti G. Joshi

**Affiliations:** Department of Biophysics, National Institute of Mental Health and Neurosciences, Hosur main road, Bangalore-560029, India; Department of Psychiatry, National Institute of Mental Health and Neurosciences, Hosur main road, Bangalore-560029, India; Department of Neuroscience, University of Virginia, Charlottesville. VA-22903, USA

**Author notes:** Correspondence: Shiksha Sharan, Ph.D.

**Keywords:** Astrocytes, Excitotoxicity, Aging, Glutamate, transporter, spinal cord.

## Abstract

Astrocytes are abundant and heterogeneous cell types in the CNS [1]. They promote neuronal health and survival and protect neurons from glutamate-induced excitotoxicity. In the spinal cord, astrocytes are present in both the dorsal horn (DH) and ventral horns (VH). However, only motor neurons in the VH are highly vulnerable to glutamate excitotoxicity. We hypothesize that the preferential vulnerability of motor neurons may underlie their spatial confinement to the VH, where astrocytes are differentially equipped with glutamate-handling machinery. With aging, glutamate excitotoxicity increases, suggesting compromised astrocytic glutamate handling. We tested our hypotheses by comparing astrocytic morphology, expression of glutamate transporters, and glutamate uptake function in the DH and VH using immunohistochemical staining, western blotting, and whole-cell patch-clamp electrophysiology. We found a global reduction in the numerical density and astroglial coverage in the spinal cord, with a prominent decline in the VH with aging. Astrocytic glutamate transporters, Excitatory amino acid transporter 1 (EAAT1) and Excitatory amino acid transporter 2 (EAAT2), show an overall reduction in expression with aging, with a more progressive decrease in the VH than DH. We performed whole-cell patch-clamp studies on DH and VH astrocytes to assess the functional outcome of the altered expression of EAATs with aging. Both VH and DH astrocytes showed a dramatic decline in glutamate uptake currents with aging, suggesting compromised glutamate handling in aging astrocytes. Our study suggests that astrocytes in the DH and VH undergo differential changes with aging, exhibiting compromised glutamate handling properties, which may be one of the key contributors to the late onset of neurodegenerative disorders.

## 1. INTRODUCTION

Astrocytes are the most abundant glial cells in the CNS [2]. They play numerous roles, including metabolic and trophic support, ion homeostasis, neurotransmitter buffering, and maintaining blood-brain barrier (BBB) integrity [3,4]. Astrocytes are the primary homeostatic cells of the central nervous system (CNS), possessing a diverse range of molecular programs and signaling pathways responsible for maintaining the health and functionality of neural tissue. Through these programs, astrocytes exert influence and regulate synaptic transmission and the functional activity of neuronal ensembles, which contribute to the overall functional output of the CNS. By actively participating in the modulation of synaptic function, astrocytes contribute to the overall balance and stability of neural circuits, ensuring optimal brain function [5]. Recent studies provide evidence of a dynamic bidirectional interaction between astrocytes and neurons that regulate neuronal activity, thereby generating behavioral patterns [6].

One of the major functions of astrocytes is to uptake the excess extracellular glutamate released during synaptic activity due to the strategic placement of their fine processes capable of efficient glutamate clearance at synapses. Astrocytic processes can be divided into three categories: branches, leaflets, and endfeet, based on their distinct morphological and functional properties. These divisions allow for specialized interactions and functions of astrocytes in the brain. Astrocytic processes are distributed among synapses, with individual synapses making contact with either branches or leaflets, or sometimes both [7].

Spines, which are small protrusions on dendrites, contain neurotransmitter receptors. On the other hand, the leaflets of astrocytic processes are responsible for hosting glutamate transporters, Na+/Ca2+ exchanger (NCX), Na+/K+ ATPase (NKA), and K+ channels. Additionally, the leaflets also house neurotransmitter receptors and other SLC (solute carrier) transporters. These specialized components within astrocytic processes enable them to regulate neurotransmitter levels, maintain ion homeostasis, and participate in synaptic signaling [7,8].

Although glutamate is the major excitatory neurotransmitter responsible for normal CNS functioning, an abnormally high level of glutamate poses risks such as excitotoxicity, oxidative stress, neuronal hyperexcitability, and neuronal death [9]. Astrocytes play important roles in maintaining extracellular glutamate levels [10]. A combination of 5 Excitatory Amino Acid Transporters (EAAT1-5), expressed strategically, acts in concert to remove excess glutamate with high efficiency [11,12]. EAAT1/GLAST and EAAT2/GLT1 are located primarily on astrocytes, and EAAT2 alone accounts for approximately 90% of all glutamate transport [13,14,10]. Although EAAT2 is believed to be the major transporter subtype in removing excess glutamate from the synaptic cleft, EAAT1 also plays a critical role in preventing excitotoxic neuronal injury [15]. EAAT3 is predominantly expressed by neurons [16]. EAAT4 is a neuron-specific glutamate transporter specifically expressed by Purkinje cells, and EAAT5 is selectively expressed in the retina [17]. Astroglial dysfunction due to altered EAAT expression contributes to glutamate-induced excitotoxicity and has been implicated in many neurological diseases, including seizures and Amyotrophic Lateral Sclerosis (ALS), as well as during normal aging [18,19,20,21,22,23,24].

Aging is a key contributor to the progression and aggravation of neurodegenerative diseases like ALS, increasing the risk factor for neurodegeneration and worsening the severity of the disease [25, 26, 27]. It has been observed that aged individuals experience more severe forms of the disease and incomplete recovery after damage [28].

The hypertrophy of astrocytes has long been recognized as a widespread indication of central nervous system (CNS) pathology. Reactive astrocytes, which undergo hypertrophy in response to pathological conditions, may exhibit maladaptive behaviors and become dysfunctional. These dysfunctional astrocytes not only lose their homeostatic functions but can also acquire detrimental functions, further worsening the existing pathology [29].

In aged human brains, there are generally mild and diverse alterations in astrocyte morphology or GFAP levels. Studies conducted on rodents have documented region-specific and sometimes conflicting changes in aging astrocytes, including increased cellular volume and overlapping processes, as well as atrophy, elevated GFAP content, or even a decrease in the number of astrocytes positive for GFAP and GS markers [29, 30].

With the normal aging process, a decline in physiological function, increased neuroinflammation, brain shrinkage, and memory deficits have been reported [31]. Astrocytes have been found to play a role in the heightened inflammatory state of the aged central nervous system (CNS). The age-related changes in astroglial morphology result in an increased presence of hypertrophic and reactive astrocytes in the aged brain.

The territorial domains of astrocytes in aged mice were found to be approximately 30% smaller compared to adult animals. Analysis of volume fraction measurements demonstrated significant dystrophy of the peripheral leaflet-like processes, which are important for astrocytic synaptic coverage. However, it is worth noting that the number of astrocytes remained unchanged in the aged mice compared to the adult animals [32, 33].

Astrocytes obtained from the aged cortex demonstrate heightened production of IL-6 compared to those from the young cortex. Additionally, the elevated levels of TNF-α, IL-1β, and IL-6 found in the aged cortex and striatum are primarily localized with astrocytes, rather than microglia or neurons. These findings suggest that astrocytes play a significant role in creating an inflammatory environment within the aged central nervous system (CNS) [29]. Moreover, molecular and cellular changes occurring during aging render neurons more susceptible to excitotoxicity, a phenomenon linked to neurodegenerative disorders as supported by studies conducted on mouse models.

Several studies have demonstrated that aged mouse cortical astrocytes exhibit upregulation of genes associated with immune responses, while the expression of glial fibrillary acidic protein (GFAP) and genes involved in neuroprotection and neuronal support is decreased. Researchers have also observed age-dependent shrinkage in astroglial peripheral processes [34]. Aging is characterized by intricate and region-specific changes in both gene expression and the structure of astrocytes. Notably, certain brain regions, such as the hippocampus, exhibit an increase in GFAP expression and alterations in the astroglial cytoskeleton. Furthermore, aging is generally accompanied by an elevated ratio of glutamate to glutamine in the brain, suggesting potential abnormalities in the functioning of the glutamate (GABA)-glutamine shuttle. Additionally, the expression of astrocytic glutamate transporters and the efficiency of glutamate uptake are reduced in aged rats (24–27 months old) compared to young adults (3–5 months old). These findings collectively contribute to our understanding of the changes that occur in aged astrocytes and their potential implications in aging-related processes [35,34]

With aging, astrocytes also undergo changes that significantly reduce their ability to support neuronal functions in the brain as well as in the spinal cord [36]. Since most neurodegenerative diseases have a late onset, and neurons are particularly susceptible to glutamate excitotoxicity, which can result from dysfunctional glutamate homeostasis [37, 38, 39], it is crucial to understand the glutamate handling properties of astrocytes in relation to aging [21].

One of the emerging attributes of astrocytes in recent decades is their structural and functional heterogeneity, suggesting that astrocytes adapt according to the specific environment in which they exist [40]. For example, astrocytes in the ventral horn express higher levels of inward rectifying K+ channel (Kir 4.1) compared to those in the DH [41]. Recent research has revealed distinct expression patterns of astrocytic homeostatic proteins, including Kir4.1, around alpha motor neurons in the ventral horn [41, 42]. Additionally, impaired potassium (K+) and glutamate homeostasis by astrocytes have been implicated in the pathology of various diseases such as amyotrophic lateral sclerosis (ALS) and Huntington’s disease [43, 44]. These findings highlight the importance of astrocytes in maintaining proper K+ and glutamate levels and suggest their involvement in the pathogenesis of certain neurodegenerative conditions. The excitability properties of DH neurons in aged animals exhibit differences compared to recordings from tissue prepared from young animals [45]. Additionally, in diseases such as ALS, motor neurons in the ventral horn are preferentially susceptible, sparing other spinal sensory neurons, as shown by several studies, including ours [46, 47, 48, 49, 50].

A large number of studies have established that spinal sensory and motor neurons residing in the DH and VH, respectively, not only differ morphologically but also functionally. However, evidence of such regional differences in astrocytes and how aging alters astrocytic properties is less explored. In the present study, we investigated whether the properties of astrocytes surrounding motor neurons differ from those surrounding sensory neurons and if this regional difference in astrocytic properties makes motor neurons susceptible to glutamate excitotoxicity. We also examined astrocytic properties in three distinct age groups to address changes associated with normal aging. The study utilizes the spinal cord as a model system to examine the astrocytic environment surrounding highly vulnerable motor neurons in the ventral horn and spatially distinct, less susceptible sensory neurons in the DH, enabling the investigation of these environments separately and without overlap.

## 2. RESULTS

### 2.1. Distinct density and morphology of astrocytes in DH and VH of rat spinal cord and changes with age

In the spinal cord, astrocytes ensheath motor and sensory neurons in the VH and DH, respectively. However, only motor neurons are highly vulnerable to glutamate excitotoxicity, and this vulnerability gradually increases with age [25, 27]. Since astrocytes are the primary cells responsible for clearing extracellular glutamate, the higher vulnerability of motor neurons may underlie the structural and functional differences in DH and VH astrocytes.

To investigate this, we first assessed the numerical density of astrocytes with aging and whether any differential changes were occurring between DH and VH astrocytes. Using the astrocyte-specific marker S100β in transverse spinal cord sections, we observed a remarkably lower number of astrocytes in the spinal cord with aging (Fig. 1A, B). The density of astrocytes was highest in the juvenile spinal cord (131±5 astrocytes/mm^2^; n = 5 rats), which decreased to over 50% in young adults (56±3 astrocytes/mm^2^; n = 5 rats) and old age (49±2 astrocytes/mm^2^; n = 5 rats) (Fig. 1B).

**Figure 1.**
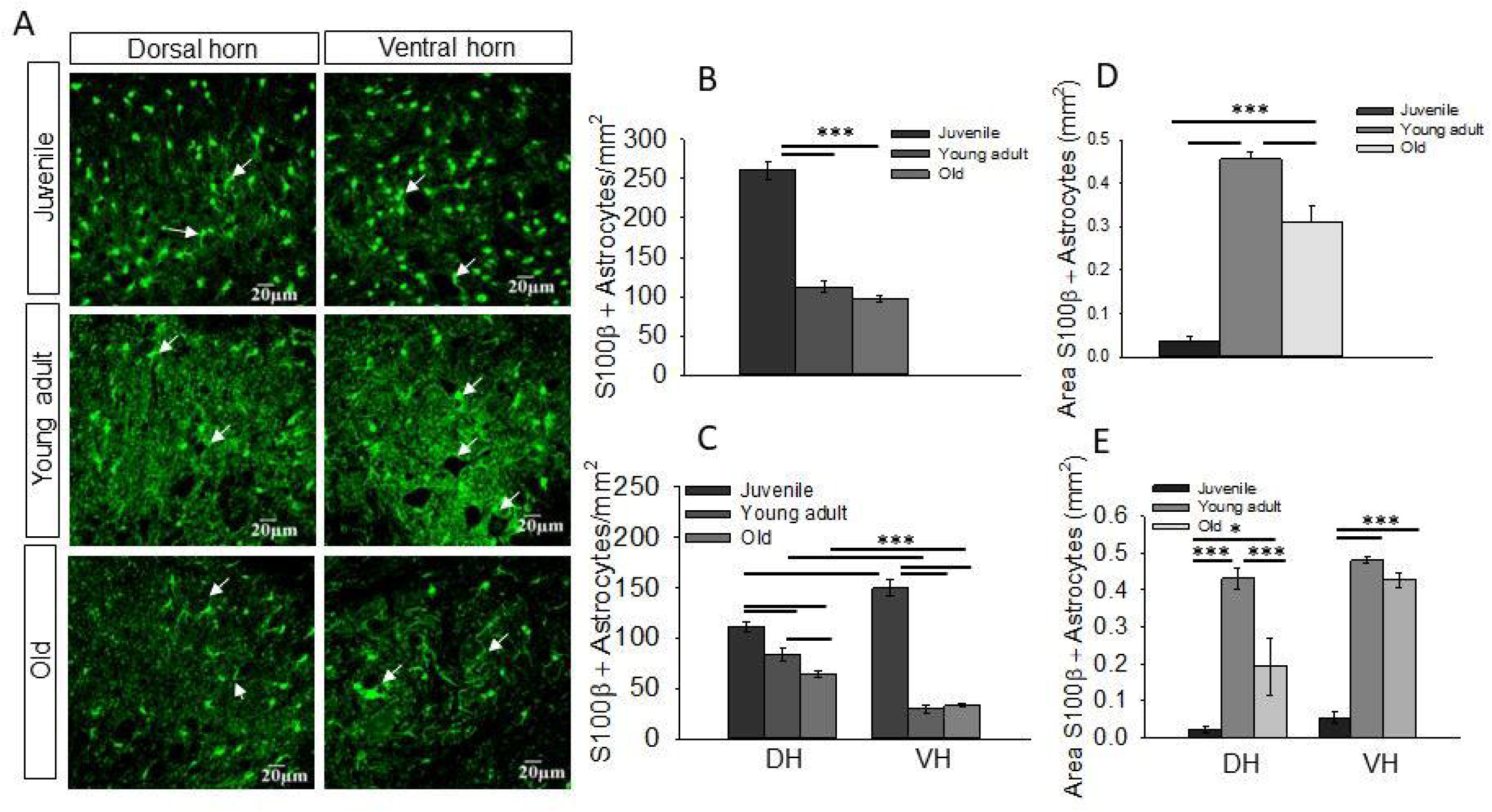
Density of astrocytes decreases with aging in of spinal cord. (A) Representative images of immunohistochemistry showing S100β -labelled astrocytes in dorsal horn and ventral horn of spinal cord, in three age groups. White arrows display astrocytes surrounding neuronal cell body. (B) the density of S100β -labelled astrocytes significantly decreased with aging in spinal cord. (C) Density of astrocytes decreased in dorsal horn and ventral horn regions of spinal cord with aging. (D) despite the number of astrocytes going down with age the area acquired by astrocytes increased with age, area acquired by young adult astrocytes was highest. (E) Graph shows same pattern in dorsal horn and ventral horn regions of spinal cord with least area acquired by juvenile astrocytes and maximum area acquired by young adult astrocytes. Data represent the mean ±S.E.M., n= 3-5 rats. (*P ≤ 0.05; **P≤0.01; ***P ≤ 0.001)

Next, to ascertain the differences in the astrocytic milieu of sensory and motor neurons with aging, we compared the density of S100β-labeled astrocytes in the DH and VH of all age groups. In juvenile rats, the VH showed a higher density of astrocytes than the DH (VH 75±4 astrocytes/mm^2^, DH 56±2 astrocytes/mm^2^; n = 5 rats) (Fig. 1A and 1C). Furthermore, in the VH of young adults (DH 41±3 astrocytes/mm^2^, VH 15±1 astrocytes/mm^2^; n = 5 rats) and old age groups (DH 32±1 astrocytes/mm^2^; VH 16±0.81 astrocytes/mm^2^; n = 5 rats), a significantly lower number of S100β- positive astrocytes were observed compared to the DH (Fig. 1C). Morphologically, astrocytic processes were less intricate in the juvenile group; however, S100β immunofluorescence in the astrocytic processes appreciably increased in the young adult and old age groups, and the processes appeared more elaborate and profusely surrounded the neurons in both the DH and VH (Fig. 1A, white arrows). This data suggests that the VH possesses a higher density of astrocytes than the DH at an early age; however, with aging, the total number of astrocytes decreases in both regions, with a more severe decline in the VH than the DH.

Since astrocytes undergo morphological maturation during development, which involves an increase in astrocytic arborizations and coverage [51], we examined whether a decline in astrocyte numbers affects overall glial coverage in the spinal cord. The juvenile age group, which exhibited a high numerical density of astrocytes (Fig. 1B), showed the lowest glial coverage (Fig. 1D, 37±9 area/ mm^2^; n = 3 rats). However, glial coverage increased approximately 48-fold in young adults (Fig. 1D, 456±17 area/ mm^2^; n = 4 rats) and subsequently decreased significantly in old age (Fig. 1D, 309±36 area/ mm^2^; n = 4 rats). In the DH and VH, astrocytic coverage displayed a similar increase from juvenile (Fig. 1E, DH 21±7 area/ mm^2^; VH 53±14 area/ mm^2^; n = 3 rats) to young adult stages (Fig. 1E, DH 430±29 area/ mm^2^; VH 481±8 area/ mm^2^; n = 4 rats), but only the DH exhibited a significant decline in the old age group (Fig. 1E, DH 193±77 area/ mm^2^; VH 426±20 area/ mm^2^; n = 4 rats). These findings suggest that although the numerical density of astrocytes decreases with age, glial coverage appears to remain stable and only decreases in the DH during old age.

### 2.2 Change in expression of astrocytic glutamate transporters EAAT1 and EAAT2 in the spinal cord with age

Next, we investigated whether astrocytes surrounding motor and sensory neurons exhibit different levels of glutamate transporter expression and whether this expression changes with aging. We began by analyzing the overall expression levels of EAAT1 and EAAT2 in the spinal cord through western blot analysis using lumbar spinal cord tissue. As EAATs have been extensively studied in hippocampal astrocytes, we used hippocampal tissue as a positive control for transporter expression. Consistent with previous reports [52], we detected protein bands corresponding to EAAT1 and EAAT2 at approximately 60 kD and 65-67 kD, respectively (Fig. 2A-B; Fig. 3A-B).

**Figure 2.**
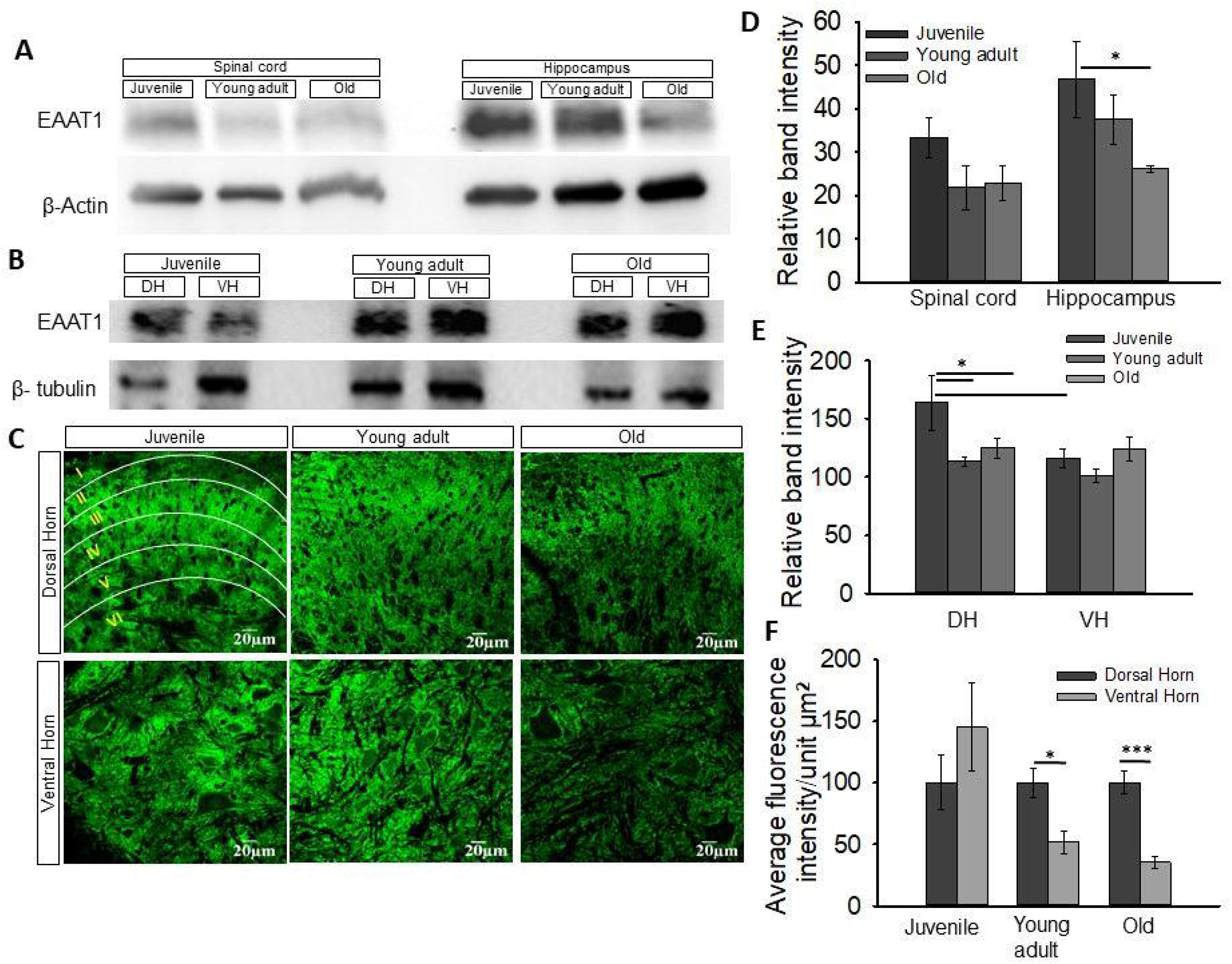
EAAT1 glutamate transporter expression decreased with aging in hippocampus of SD rat: (A -B) Representative western blots show EAAT1, and beta-actin/beta-tubulin bands. (C) Confocal image of a transverse section of rat lumbar spinal cord lamination immunostained with EAAT1 astrocytic glutamate transporter. White lines represent boundaries of the dorsal horn laminae. Astrocytes from three age groups were immunostained with EAAT1 to access effect of age in glutamate transporter expression. (D) western blot analysis showed a decline in EAAT1 expression with aging in hippocampus although there was no significant change in EAAT1 expression with age in spinal cord. (E) comparing EAAT1 expression in regions of spinal cord, EAAT1 showed decline in its expression when dorsal horn of young adult and old rats were compared with juvenile rats. Also, the expression of EAAT1 in ventral horn of juvenile rats were less than ventral horn. (F) evaluation of EAAT1 immunofluorescence in regions of spinal cord showed decreased EAAT1 expression in DH of young adult and old rats with no change in juvenile age group. Western blot data represent the mean ±S.E.M. of n=3 rats. IHC data represent the mean ±S.E.M. of n=4 rats. (*P ≤ 0.05; ** P≤0.01; ***P ≤ 0.001)

**Figure 3.**
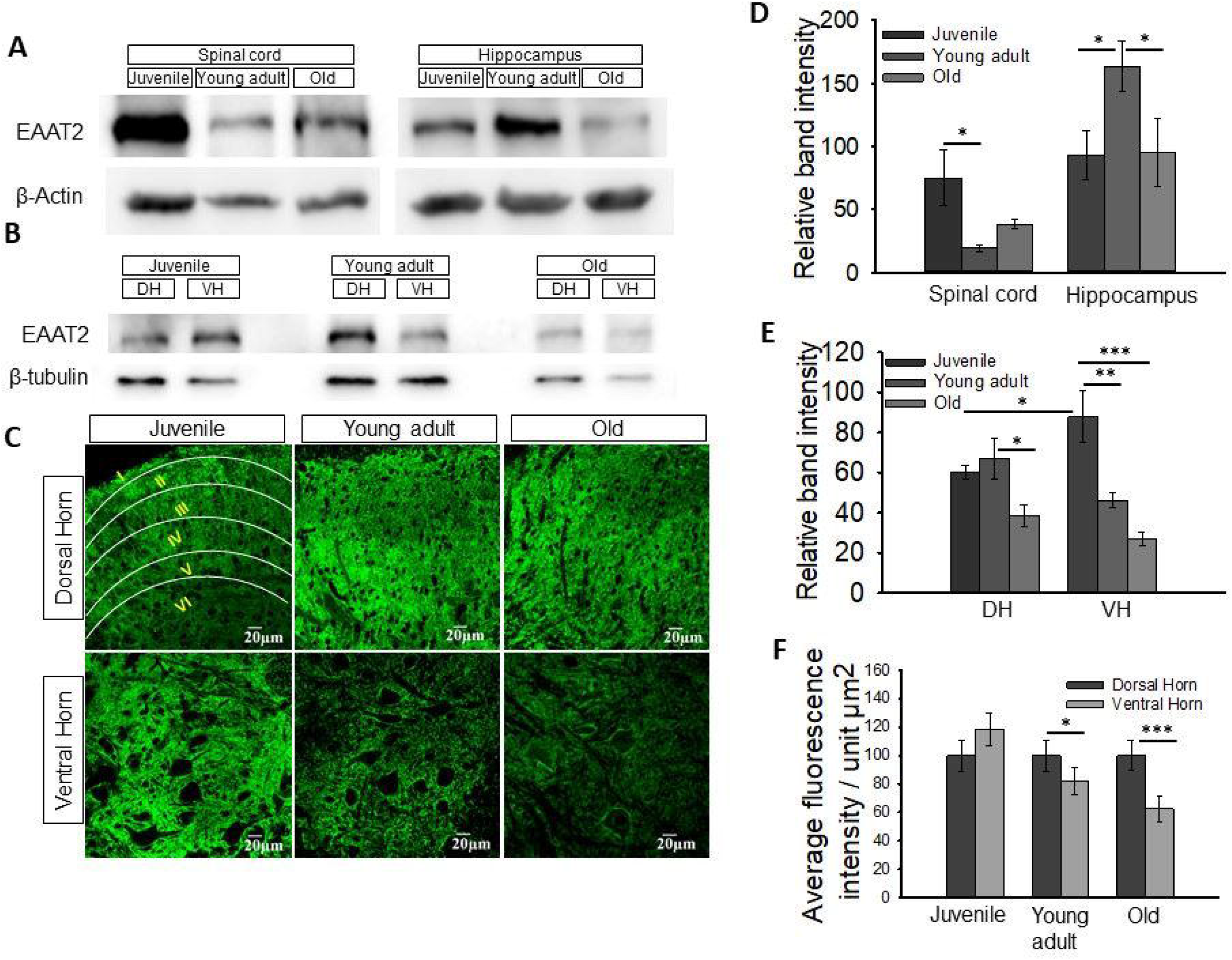
Most crucial glutamate transporter EAAT2 showed reduced expression with aging in regions of spinal cord. (A-B) western blot images of EAAT1, and beta-actin/beta tubulin bands. (C) immunostaining of astrocytes with EAAT2 transporter indicated the differential laminar expression pattern of transporter with intense expression in lamina II, III and IV of dorsal horn (white lines) especially in young adult and old groups. (D) Spinal cord tissue had highest EAAT2 expression in juvenile age group followed by old and young adult groups. However, the positive control hippocampus portrayed highest EAAT2 levels in young adult rats. (E) The expression of EAAT2 decreased with maturing in both dorsal horn and ventral horn of spinal cord with more prominent decrease in ventral horn. (F) immunofluorescence data showed lower EAAT2 expression in DH higher age groups as compared to VH astrocytes. Western blot data represent the mean ±S.E.M. of n=3 rats. IHC data represent the mean ±S.E.M. of n=4 rats. (*P ≤ 0.05; **P≤0.01; ***P ≤ 0.001)

In the spinal cord, EAAT1 expression was highest in the juvenile group (Fig. 2A, D, 33±4, n = 3 rats) and decreased with age, although not significantly, in young adults (Fig. 2A, D, 21±5, n = 3 rats) and old animals (Fig. 2A, D, 22±3, n = 3 rats). Interestingly, the positive control hippocampus also exhibited the highest EAAT1 expression in juveniles, which significantly declined with age (Fig. 2A, D, juvenile 46±8; young adult 37±5; old 26±0.8, n = 3 rats), suggesting age-related changes in EAAT1 expression independent of CNS region. Moreover, when comparing EAAT1 expression in the DH and VH, we found that the juvenile group showed significantly lower EAAT1 expression in the VH compared to the DH (Fig. 2B, E, juvenile DH 164±40; VH 116±13, n = 3 rats). As age progressed, EAAT1 expression significantly decreased in the DH, while VH remained unchanged (Fig. 2B, E, young adult DH 113±7; VH 101±9; old DH 125±14, VH 124±17, n = 3 rats). To further confirm the distribution and differential expression of EAAT1 with aging and in DH and VH, we performed immunofluorescence analysis of EAAT1 in transverse sections of the spinal cord at the lumbar level. Expression was measured as the average fluorescence intensity/unit µm2. EAAT1 immunofluorescence exhibited a distinct pattern in the DH, with robust expression in rexed lamina II/III/IV, particularly in young adult and old animals (Fig. 2C, outlined). In contrast, VH showed relatively uniform expression, although in higher-aged groups, astrocytes appeared to surround the cell bodies of motor neurons. In the young adult and old age groups, EAAT1 expression in the VH was significantly lower compared to the DH, with this difference being more prominent in old age (Fig. 2C, F, young adult DH 100±11, VH 52±9; old DH 100±9, VH 35±4, n = 4 rats).

The expression of EAAT2, which is responsible for 90% of glutamate transport in astrocytes [53], declined with age in the spinal cord. The juvenile group exhibited a 3-fold higher expression of EAAT2 (Fig. 3D, 75±21, n = 3 rats) compared to the young adult group (Fig. 3D, 19±2, n = 3 rats). However, there was no significant difference in EAAT2 expression between the old and young adult groups (Fig. 3D, old 38±4, n = 3 rats). In the positive control hippocampus, EAAT2 expression was highest in the young adult group compared to the juvenile and old groups (Fig. 3D, juvenile 93±19; young adult 163±20; old 95±27, n = 3 rats).

Furthermore, we compared the expression of EAAT2 in the DH and VH of the spinal cord. In the juvenile group, VH exhibited higher EAAT2 expression than DH (Fig. 3E, DH 60±3; VH 87±12, n = 3 rats). However, EAAT2 expression decreased in older age groups, with significantly lower expression in the VH compared to the DH (Fig. 3E, young adult DH 66±10; VH 46±3; old DH 38±5; VH 26±3, n = 3 rats). Overall, these findings indicate a clear decrease in EAAT2 levels with age, particularly in the VH. However, DH showed comparable EAAT2 expression between the juvenile and young adult groups, with old animals exhibiting the lowest EAAT2 expression (Fig. 3E).

Immunohistochemistry data revealed similar EAAT2 expression in the DH and VH of juvenile groups. However, the VH of the young adult and old groups exhibited higher EAAT2 expression compared to the DH (Fig. 3C, F, old DH 100±10; VH 62±9, n = 4 rats).

In summary, both EAAT1 and EAAT2 displayed distinct regional distribution patterns that varied to some extent among the three age groups (juvenile, young adult, and old). EAAT1 immunoreactivity was predominantly observed in lamina II compared to other laminae of the spinal cord (Fig. 2C, outlined). On the other hand, EAAT2 showed higher intensity in laminae III, IV, and V compared to laminae I and II, especially in young adult and old rats (Fig. 3C, outlined). Both EAAT1 and EAAT2 were mostly uniformly distributed in the VH of the spinal cord, with more pronounced expression surrounding motor neurons (Fig. 2C and 3C). Overall, the expression of EAAT1 and EAAT2 exhibited an overall decline in the spinal cord with aging, with VH astrocytes showing a more significant decline compared to DH astrocytes, suggesting a reduced capacity for glutamate uptake in VH astrocytes with age.

### 2.3 Glutamate uptake in DH and VH astrocytes with aging

To assess glutamate uptake by astrocytes, we measured glutamate-induced currents in astrocytes of the DH and VH in ex-vivo acute spinal cord slices, as well as in astrocytes of hippocampal slices. Astrocytes were identified and selected based on their morphology, size, location, and electrophysiological properties [54]. The resting membrane potential of astrocytes ranged from -68 mV to -70 mV (n=22), while neurons exhibited a resting membrane potential of -60 mV to -63 mV (n=3). When a 20-pA current was injected for 100 ms in the current clamp mode, neurons generated a train of action potentials, while astrocytes exhibited passive depolarization (Fig. S2). We used glutamate current recordings from hippocampal astrocytes as a positive control.

For all the glutamate uptake current recordings in astrocytes, the membrane potential was held at -70 mV, and a 10 mM glutamate pulse was applied for 1 minute using a superfusion system. Figure 4A presents representative traces of currents recorded from astrocytes in the DH, VH, and hippocampus across all three age groups. Astrocytes in the juvenile group displayed substantial glutamate uptake currents, which were comparable between DH and VH (Fig. 4B, DH 171±33 pA; VH 155±25, n=12 cells from 6 rats). We observed a significant decline in glutamate uptake currents in both DH and VH astrocytes of young adults (Fig. 4B, DH 15±4 pA; VH 13±5, n=5 cells from 5 rats) and old groups (Fig. 4B, DH 10±5 pA; VH 10±3, n=5 cells from 5 rats).

**Figure 4.**
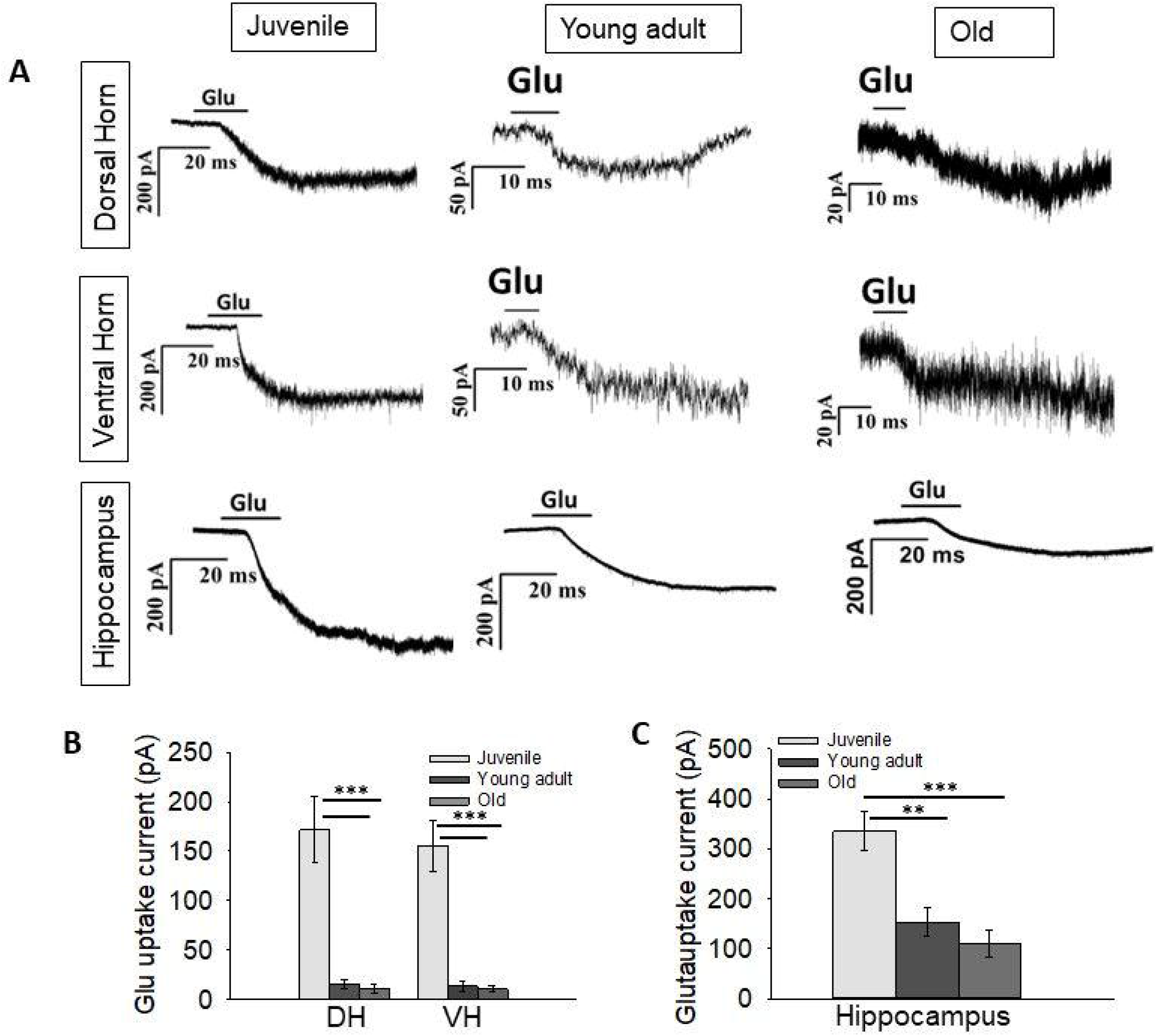
Glutamate transporter current in hippocampal and spinal astrocytes: (A) Representative astrocytic glutamate uptake current traces in spinal cord and hippocampal astrocytes with age. (B) Bar graph shows huge decline in average glutamate uptake current recorded from astrocytes of dorsal and ventral horn in spinal cord of young adult and old groups. (C) hippocampal astrocytes demonstrated decline in glutamate uptake current with growing age. [Hippocampal astrocytes (n=5-8); 15-30 Days Dorsal horn (n=12); Ventral horn (n=12); 2-3 months and 18-20 months Dorsal horn (n=5); Ventral horn (n=5). Data represented as the mean ±S.E.M. (*P ≤ 0.05; **P≤0.01; ***P ≤ 0.001)

To serve as age-matched positive controls, we also measured glutamate uptake currents in hippocampal astrocytes from the CA1 region across all three age groups. Previous studies have reported a decline in astrocytic glutamate uptake currents during maturation [35]. Our data confirmed a substantial decrease in astrocytic glutamate uptake currents with aging in the hippocampus (Fig. 4C, juvenile 335±39 pA, n=6 cells; young adult 153±28 pA, n=4 cells; old 110±26 pA, n=6 cells, from 4 rats). However, the glutamate uptake currents recorded in hippocampal astrocytes were much higher than those in spinal cord astrocytes. In the hippocampus, compared to the juvenile group, the higher age groups exhibited approximately a 67% decrease in glutamate uptake currents. In contrast, in the spinal cord, the glutamate uptake currents declined drastically by 93% in the higher age groups, with comparable uptake currents in young adult and old rats

## 3. MATERIALS AND METHODS

### 3.1 Materials

Polyclonal antibodies against EAAT1 and EAAT2 were obtained from Alomone Laboratories in Jerusalem, Israel. Secondary antibodies used were Alexa Fluor 488 goat anti-rabbit IgG, Alexa Fluor 633 goat anti-rabbit IgG, Alexa Fluor 633 goat anti-mouse IgG, and Alexa Fluor 488 goat anti-mouse IgG, all purchased from Molecular Probes, Invitrogen, Life Technologies in India. Neurochrom-cy3 conjugated, anti-GFAP-488 conjugated, Guinea pig anti-EAAT2, FITC goat anti-Guinea pig, and FITC rabbit anti-goat antibodies were obtained from Merck, Millipore India. Horseradish peroxidase (HRP) conjugated anti-rabbit and anti-mouse antibodies were obtained from Sigma Aldrich. L-Glutamate was also purchased from Sigma Aldrich. S100β was sourced from Santacruz Biotechnology. For western blotting, antibodies against EAAT1, β-actin, and tubulin were obtained from Abcam. All other chemicals used were of analytical grade and were obtained from commercial sources.

### 3.2 Experimental Animals

Sprague-Dawley rats from three age groups, namely juvenile (21 days), young adult (3 months), and old (20 months), were obtained from the Central Animal Research Facility of the National Institute of Mental Health and Neurosciences in Bangalore, India. All animal procedures were conducted in accordance with the guidelines of the institutional animal ethics committee. Briefly, rats were anesthetized using diethyl ether and subsequently decapitated.

### 3.3 Western blotting

Whole-cell membrane fractions were prepared from the DH and VH of spinal cord tissue (lumbar region) isolated from different experimental groups. In brief, rats from different age groups were anesthetized and decapitated, and the spinal cord was quickly extracted using hydraulic extraction. An 11-gauge needle with a blunted tip, filled with Artificial CSF (aCSF), was inserted into the spinal canal near the L5 segment to a depth of 1 cm. The aCSF was forcefully injected to flush out the spinal cord. The cauda equina was identified, and the alignment of the spinal cord was adjusted accordingly, taking into account the rostral and caudal ends. The lumbar section of the spinal cord was isolated and placed in ice-cold HEPES buffer for 1-2 minutes. Subsequently, the spinal cord was sliced to a thickness of 800 µm, and the DH was separated from the VH by making a horizontal slice at the level of the central canal, using a stereomicroscope. The separated DH and VH were manually homogenized using a motor and piston homogenizer in lysis buffer [containing 10 mM Tris, 5 mM KCl, 1 mM DTT, 1 mM EGTA, and Complete Protease Inhibitor Cocktail (Abcam)], and the resulting cell lysate was sonicated.

The protein concentration was determined using the Lowry method [55], and the samples were then diluted 1:1 in sample treatment buffer [containing 25 mM Tris, 8% SDS, 40% Glycerol, 0.05% Bromophenol Blue, and 20% β-Mercaptoethanol]. After denaturation by heating at 95°C for 3-5 minutes, 100 µg of protein was loaded into each lane of a 10% SDS-PAGE gel. Equal concentrations of proteins from both regions were loaded in each well and subjected to electrophoresis. Nonspecific binding sites were blocked with 5% non-fat milk, and the blots were incubated for 3 hours at room temperature with rabbit polyclonal anti-EAAT1/anti-EAAT2 (1:200) and β-actin/tubulin (1:10,000). The blots were then probed with horseradish peroxidase-conjugated anti-rabbit (1:250) or anti-mouse (1:500) secondary antibodies using Super Signal® West Pico chemiluminescent substrate. The enhanced chemiluminescence signal was captured using a camera attached to a computer-assisted gel documentation system (Syngene, U.K). Quantification of the band intensities was performed using Image J software (NIH, USA).

### 3.4 Immunohistochemistry

Immunohistochemistry (IHC) was performed on free-floating spinal cord sections using the following primary antibodies: rabbit anti-EAAT1 (1:50), Guinea pig anti-EAAT2 (1:1000), mouse anti-GFAP (1:400), and mouse anti-S100β (1:100). Transcranial perfusion was carried out with chilled 0.9% saline, followed by 4% paraformaldehyde in PBS. The lumbar region of the spinal cord was extracted and kept in a 4°C paraformaldehyde solution for 48 hours, followed by transfer to PBS for an additional 24 hours at 4°C. Sections with a thickness of 70 µm were prepared using a vibratome (Vibratome, 3000 Sectioning System). Antigen retrieval was performed by placing the sections in a solution of saline sodium citrate and formamide (1:1, volume ratio) with a pH of 6.4 for 2 hours at 65°C, using gentle shaking in a water bath. Non-specific antigen blocking and permeabilization were carried out by incubating the sections in PBS containing 5% bovine serum albumin, 5% goat serum, and 0.1% Triton X-100 at room temperature for 2 hours. The sections were rinsed with PBS and then incubated in 100 µl of PBS containing the primary antibody, 5% goat serum, and 5% bovine serum albumin for 12 hours at 4°C. After rinsing 5 times with PBS, Alexa fluor-488/633 conjugated or FITC conjugated secondary antibodies were added and incubated for 6 hours in the dark at room temperature. The sections were thoroughly rinsed with PBS and mounted on gelatin-coated slides using a mounting solution consisting of 50% glycerol and 50% PBS (volume ratio). Images were acquired using a laser scanning confocal microscope (Olympus FV 1000, Olympus Japan).

### 3.5 Quantification of receptor expression

For the quantification of receptor expression in immunofluorescence analysis of EAAT1 and EAAT2, images were acquired from the DH and VH of all three age groups. A minimum of 3 spinal cord sections from 3 different animals in each age group were selected for analysis. Image acquisition parameters were kept consistent for all age groups. Uniform-sized regions of interest (ROIs) were drawn in specific spinal cord areas of all the groups, and the average fluorescence intensity was obtained using Fluo-view software (Olympus, Japan). The VH percentages were calculated relative to the DH, which was normalized to 100% [56]. Data representing the mean ± SEM were compared among the different experimental groups.

### 3.6 Quantification of astrocytic numerical density and coverage

For the quantification of astrocytic numerical density and coverage, S100β-labeled spinal cord sections were used. The distribution of astrocytes in the DH, VH, and the entire spinal cord was analyzed and compared across different age groups. Cell counting and area analysis were performed on the acquired confocal microscope images using ImageJ NIH image analysis software (http://rsb.info.nih.gov/ij/download.html). For cell counting, the acquired images were converted into 8-bit binary images, and a threshold was set. The cells were then counted using an overlay of the original and binary images. For astrocytic coverage, the total area occupied by astrocytes was calculated using an automated binary thresholding method in ImageJ. The area covered by cells was measured and analyzed.

### 3.7 Acute slice preparation for whole-cell patch-clamp recording

Transverse slices from the lumbar region of the spinal cord and coronal brain slices (300 µm) enclosing the hippocampus were prepared using a vibratome (Vibratome, 3000 Sectioning System) from deeply anesthetized Sprague-Dawley rats of specific age groups. The brain/spinal cord was swiftly removed and placed in an ice-cold cutting buffer containing the following concentrations (in mM): 215 Sucrose, 2.5 KCl, 1.25 NaH2PO4, 0.5 CaCl2, 4 MgSO4.7H2O, 26.2 NaHCO3, 1 Ascorbate, equilibrated with a mixture of 95% O2 and 5% CO2. The tissue was affixed to the slicing stage of a Leica vibratome. Subsequently, the slices were incubated in oxygenated minimum essential medium eagle (MEM) at room temperature (RT) for 1 hour. Whole-cell patch-clamp recordings were then conducted at RT using pipettes with resistances typically ranging from 6-8 MΩ (P-2000, Sutter Instruments, USA). The pipette solution consisted of the following concentrations (in mM): 145 potassium gluconate, 5 NaCl, 2 MgCl2, 10 HEPES, 0.2 EGTA. Astrocytes were visually identified and selected based on their size, location, and electrophysiological properties using infrared optics on an upright microscope (Eclipse E600-FN Nikon) (Fig. S1). Inward currents were recorded at -70 mV [41] using Patchmaster software, EPC10 amplifier, HEKA, Germany. The recorded data were filtered at 10 kHz (Bessel filter) and compensated for C-fast and C-slow. L-glutamate (10 mM) was applied through superfusion without altering the perfusion rate and temperature.

### 3.8 Statistical Analysis

For statistical analysis, the OriginPro or SigmaPlot programs were employed, and the data are presented as mean ± SEM. In experiments with two groups, an unpaired t-test was used to determine significance, while experiments with more than two groups were assessed using One-Way ANOVA followed by the Fisher LSD posthoc test. The threshold for statistical significance was set at p ≤ 0.05. In the figures, * represents p ≤ 0.05, ** represents p ≤ 0.01, and *** represents p ≤ 0.001.

## 4. DISCUSSION

It is well-known that neuronal degeneration occurs during normal aging, and the onset of neurodegenerative disorders is typically associated with aging [25, 57]. Glutamate-induced excitotoxicity has been implicated not only in the degeneration of neurons associated with aging but also in various neurodegenerative disorders. Recent studies have revealed that astrocytes undergo senescence and play a role in the aging process as well as in disease development [58, 59]. Das and Svendsen (2015) reported that astrocytes derived from the spinal cords of aged rats and SOD193A mutant rats exhibit senescence phenotypes and have a reduced capacity to support motor neurons [36]. These studies were performed in astrocyte cultures derived from rat spinal cords, and their impact on the survival of co-cultured NSC34 motor neuron cells was assessed. However, most studies conducted in this field do not accurately replicate the neuron-astrocyte interaction as it occurs in vivo, and there is a significant lack of in vivo research in this area. In our study, our aim was to investigate the impact of aging on astrocyte properties in an in vivo setting, with a specific focus on the neuron-astrocyte interaction in the spinal cord. We examined the characteristics of astrocytes located in the VH and DH of the spinal cord, with particular emphasis on glutamate transporters.

Additionally, we examined the influence of aging on astrocyte characteristics. Our data demonstrate age-related changes in astrocytes that are considered unfavourable for neuronal survival, and these changes are generally more pronounced in the astrocytes surrounding motor neurons in the VH of the spinal cord. In the older age group, lumbar spinal cord slices showed a decreased density of S100β-labeled astrocytes, suggesting a decline in astrocyte density with age. These changes were particularly prominent in the VH. Area analysis of S100β-positive astrocytes revealed that astrocytic coverage increases with age, which could be attributed to age-related mild astrocytic hypertrophy without concurrent pathology [60]. The hypertrophy observed in aging does not mimic the astrogliosis seen in the young brain following injury [61, 62, 63]. Upon activation, astrocytes secrete factors that modulate neuronal survival, with activated astrocytes producing neurotrophic factors that promote neuronal growth [64].

Many studies have been conducted on the expression and function of glutamate transporters in the brain, but limited information is available regarding glutamate transport in the spinal cord. Furuta et al. (1997) investigated the expression of glutamate transporters in the central nervous system (CNS) during development and demonstrated that the expression of glutamate transporter subtypes changes during development, from embryonic day 15 to postnatal day 24 [65]. However, there are no reports on the expression of glutamate transporter proteins and their activity in the spinal cord of aged animals. Our data reveal a decline in glutamate transporter expression with aging in spinal cord astrocytes, particularly in the VH. The glial glutamate transporters EAAT1 and EAAT2 were observed to be expressed in both regions of the spinal cord in all age groups, but their expression significantly declined with aging. Additionally, the expression of both transporters was notably lower in the VH compared to the DH.

We measured glutamate-induced currents in astrocytes using the whole-cell patch clamp technique. Astrocytes in the juvenile group exhibited substantial glutamate uptake currents, albeit significantly lower than the glutamate uptake currents observed in the age-matched hippocampus. The glutamate uptake current was suppressed with aging, both in the hippocampus and the spinal cord. A decrease in the expression of glial glutamate transporters EAAT1 and EAAT2, as well as reduced glutamate uptake, was observed in the aged Sprague Dawley rat hippocampus. Similar findings have been reported in the striatum of the hippocampus, where glutamate clearance was compromised in the aged brain [52]. These differences in glutamate uptake currents may reflect either varying demands on glutamate homeostasis or compromised glutamate handling capability with aging. Extensive evidence suggests that glutamate uptake into astrocytes occurs through sodium-dependent excitatory amino acid transporters (EAATs), and studies have shown that the majority of glutamate uptake is mediated by astrocytic glutamate transporter EAAT2 [13, 10]. Interestingly, the decline in glutamate uptake current was more pronounced in the spinal cord. Compared to the juvenile group, the uptake current was at least ten times lower in young adult and older rats (Fig. 4). Nonetheless, appreciable glutamate transporter expression was observed in the old age group. Several studies have demonstrated that an increase in intracellular glutamate and sodium ions leads to a decrease in glutamate transporter currents in astrocytes [66]. The activity of glutamine synthase (GS) in astrocytes dynamically regulates intracellular glutamate concentration [67], thus influencing glutamate transport across the membrane. Souza et al. (2015) reported a decrease in astrocytic GS activity in adult and old rats [68]. However, we measured glutamate uptake currents under whole-cell voltage clamp conditions, and the intracellular glutamate concentrations under the given experimental conditions would be similar in all age groups. Therefore, the observed decline in glutamate uptake current in the old age group is less likely to be attributed to the factors mentioned above.

In essence, our study provides evidence of impaired glutamate management by astrocytes with aging. The crucial role of astrocytes in maintaining extracellular glutamate levels at the synaptic cleft is well-established. The present study indicates compromised glutamate uptake capacity of astrocytes with aging, as both the expression of transporters and glutamate uptake capacity decline with age. Additionally, we report differential morphological and functional properties of DH and VH spinal astrocytes. Since we have observed a decrease in astrocytic glutamate transporter expression with aging, especially in the VH, it will be interesting to elucidate.

## Abbreviations

DH: Dorsal Horn
VH: Ventral Horn
CA1: Cornu Ammonis 1
EAAT1: Excitatory Amino Acid Transporter 1
EAAT2: Excitatory Amino Acid Transporter 1

## Author’s contribution

Shiksha Sharan: Conceptualization, conducted experiments, methodology, data curation, visualization, Investigation, validation, writing an original draft and reviewing; Bhanu Prakash Tewari: Methodology, writing, reviewing, editing; Preeti G. Joshi: supervision, conceptualization, visualization, methodology, validation.

## Acknowledgments

Animal facility: Central Animal Research Facility, National Institute of Mental Health & Neurosciences, Bangalore, India. Funding: SS received a fellowship from the Department of Science and Technology, Government of India.

---

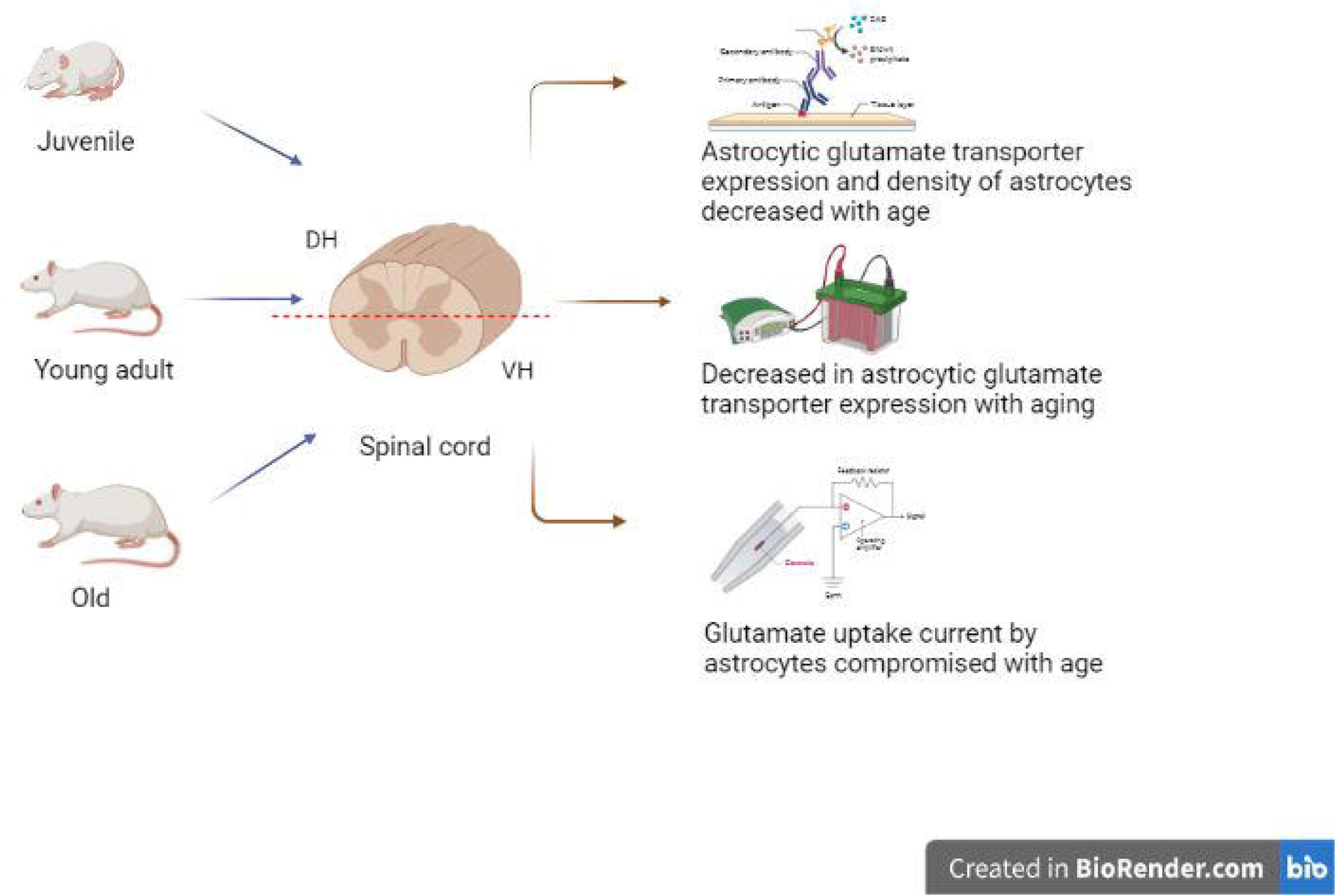

