## Supplementary material for "Aging-related changes in expression and function of glutamate transporters in rat spinal cord astrocytes": Figure S1 and S2: legends S.rtf

Figure S1. An image of a dye filled patched astrocyte (Alexa 488 channel, 20-30 ìm deep in the slice, single optical section. 

Figure S2. Comparison of electrophysiological properties of neurons and astrocytes in rat spinal cord slice. The amplitudes of the membrane potentials are plotted against 20 pA injected current step for 400 ms. Traces show electrical response of a (A) neuron and (B) astrocyte. neurons upon current injection fired action potential, whereas astrocytes were depolarised.
