## Supplementary figures and images for "Aging-related changes in expression and function of glutamate transporters in rat spinal cord astrocytes"

### Figure S1.tiff

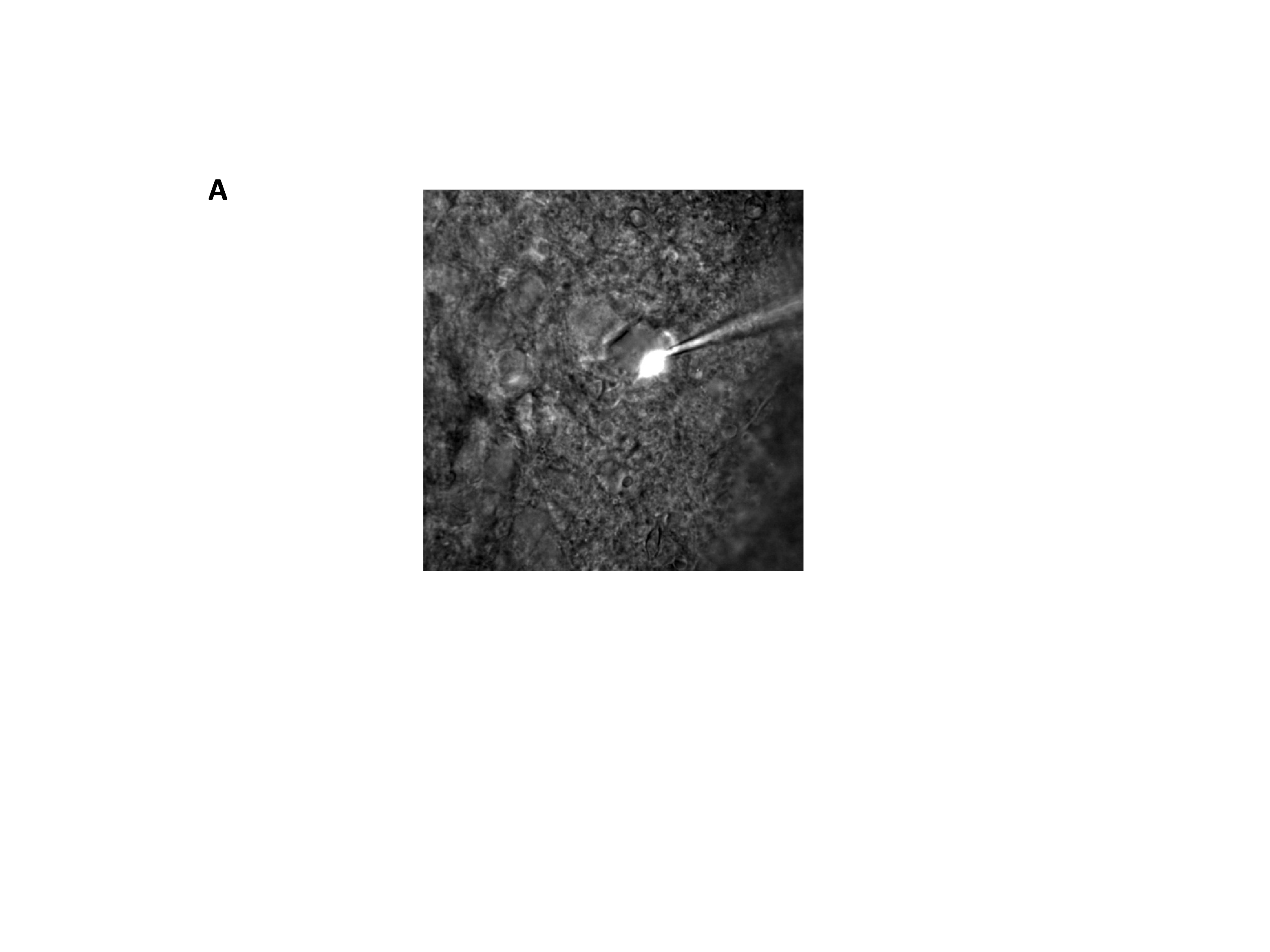

### Figure S2.tiff

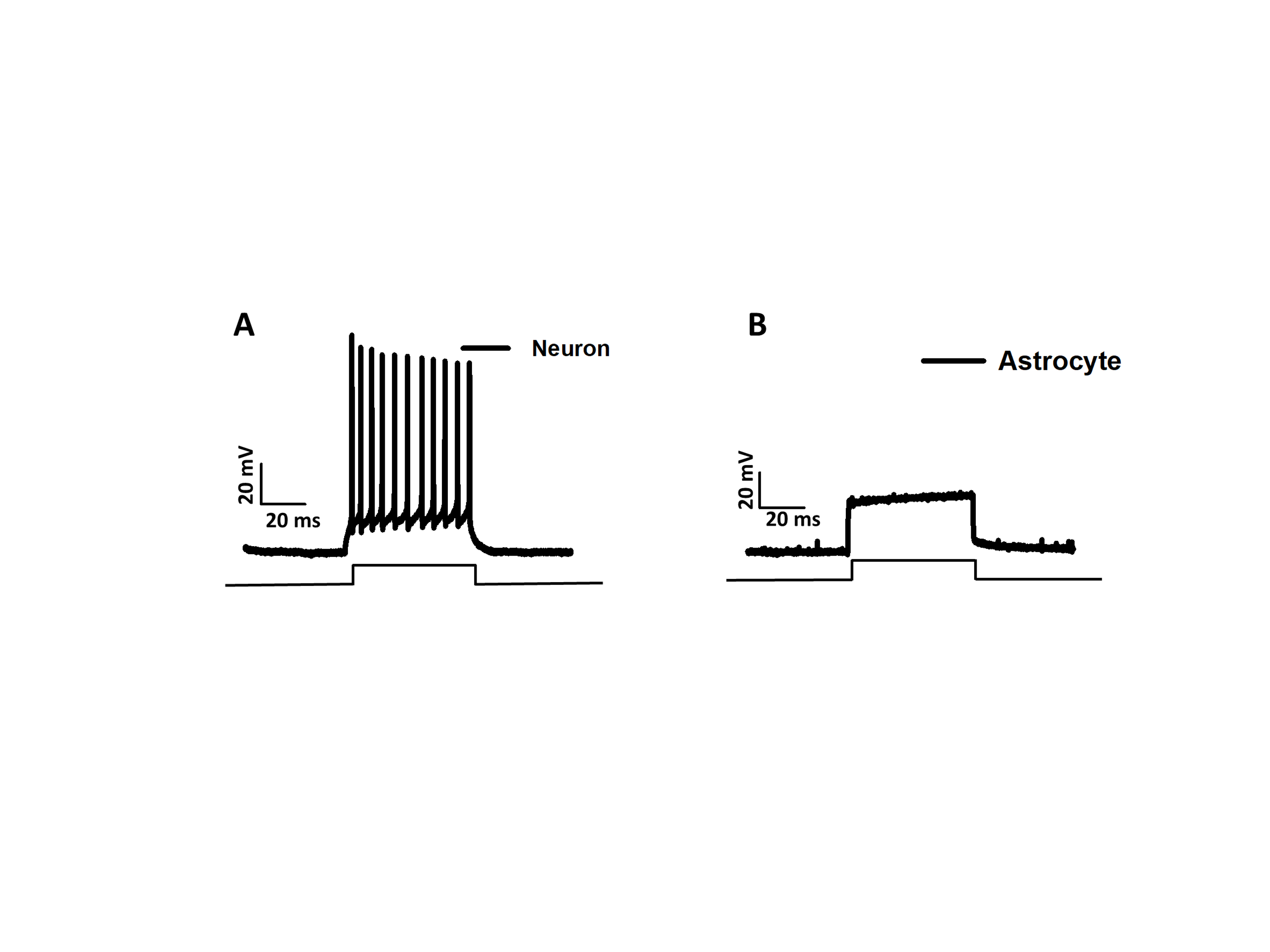
